# Thin-diaPASEF: diaPASEF for maximizing proteome coverage in single-shot proteomics

**DOI:** 10.1101/2024.04.26.591246

**Authors:** Ryo Konno, Masaki Ishikawa, Daisuke Nakajima, Kaori Inukai, Osamu Ohara, Yusuke Kawashima

**Affiliations:** Department of Applied Genomics, Kazusa DNA Research Institute, Kisarazu, Chiba 292-0818, Japan; Department of Pediatrics, Tokai University School of Medicine, 143 Shimokasuya, Isehara, Kanagawa 259-1193, Japan; Kazusa Genome Technologies Inc., Kisarazu, Chiba 292-0818, Japan; Graduate school of science, Kitasato University, Sagamihara, Kanagawa 252-0373, Japan

**Keywords:** timsTOF HT, diaPASEF/Thin-diaPASEF, Deep Proteomics, Mass Spectrometry (MS), HEK Cells, Biomarker Discovery

## Abstract

Proteomics using mass spectrometry (MS) has significantly advanced, offering deep insights into complex proteomes. The timsTOF MS platform with its parallel accumulation-serial fragmentation (PASEF) technology has achieved high scan speeds and high-quality spectra. Buker’s timsTOF HT, which features TIMS-XR technology, offers an improved dynamic range and analysis depth, supporting high sample loadings. Moreover, various improvements to the data-independent acquisition method based on the PASEF technology (diaPASEF) have been reported. Despite these advancements, most high-level deep proteomic reports are based on the Orbitrap Astral and Orbitrap Exploris 480, and analytical systems using timsTOF MS still require improvement. Here, Bruker’s timsTOF HT was used to validate and optimize key diaPASEF parameters, leading to the development of a Thin-diaPASEF method. This method provides a high quantitative accuracy and consistency. In our validation, 9,400 proteins were identified in a single shot from HEK cells (strictly controlled Protein false discovery rate < 1%), the highest number analyzed by the timsTOF MS series using standard human cultured cells. Furthermore, by combining Thin-diaPASEF with an improved *Lycopersicon esculentum* lectin method, over 5,000 proteins were identified in a 24-sample/d analysis, and we succeeded in constructing a system with high proteome coverage that can be used for biomarker discovery.

## Introduction

Proteomics using mass spectrometry (MS) has evolved rapidly as a key discipline for uncovering molecular mechanisms underlying biological processes. Recent advances in mass spectrometry and improvements in data-independent acquisition (DIA) have enabled deep proteomics^1–3^ to offer unparalleled insights into complex proteomes. As proteomics continues to expand to more complicated and clinically relevant samples, there is an increasing demand for methods with high proteome coverage and precise quantification.^4^

The timsTOF MS platform is distinguished by its dual-trapped ion mobility spectrometry (TIMS) device, in which one TIMS unit traps and concentrates ions, whereas the other performs high-resolution ion mobility separation. The system achieves an unprecedented scan speed by integrating Dual TIMS devices with parallel accumulation-serial fragmentation (PASEF) technology. Furthermore, a DIA method based on PASEF technology (diaPASEF) has been established,^5,6^ enabling analysis with high-quality spectra and reduced spectral complexity due to ion mobility. Recently, new diaPASEF methods, such as slice-PASEF,^7^ midia-PASEF,^8^ and synchro-PASEF,^9^ have been developed, which offer improved performance.

Buker’s timsTOF HT, with its 4th-generation TIMS-XR technology and advanced digitizer,^10^ offers an improved dynamic range and greater depth of analysis. Enhanced TIMS-XR technology allows for high sample loadings, which are crucial for maximizing the number of identified proteins. While earlier TIMS models also allowed for increased ramp times, the enhanced dynamic range and digitizer technology of the timsTOF HT make this a more effective strategy for ion enrichment, enabling higher sample loadings and greater flexibility in optimizing diaPASEF parameters, including sample loading and ramp time adjustments.

Although Buker’s timsTOF MS series has improved MS technologies and MS data acquisition methods, deep proteome analysis that detects over 10,000 proteins in a single-shot analysis is mostly performed using Thermo Fisher Scientific instruments, such as the Orbitrap Exploris 480 MS and the very expensive Orbitrap Astral MS.

In single-shot analysis, over 10,000 proteins were reported to be detected from standard HeLa and HEK cells using Thermo Fisher Scientific’s instruments,^3,11,12^, while a maximum of approximately 8,962 proteins were reported to be detected from HeLa cells using Buker’s timsTOF MS series^13^; previously, it was found that Buker’ MS instruments were significantly inferior to Thermo Fisher Scientific’ MS instruments in deep proteome analysis.

In this study, we used timsTOF HT to systematically validate and optimize key diaPASEF parameters, including polygon area, isolation window width, ramp time, and accumulation time for deep proteomics. We named this diaPASEF method thin-diaPASEF. Thin-diaPASEF provides higher quantitative accuracy than other diaPASEF methods while ensuring consistency and reproducibility. By combining Thin-diaPASEF with an improved version of our previously developed plasma pretreatment method, the *Lycopersicon esculentum* lectin (LEL) method,^14^ we successfully identified over 5,000 proteins in 24-sample/d analysis. Thin-diaPASEF can provide comprehensive proteomic coverage and facilitate biomarker discovery.

## Experimental procedures

### Ethics approval and consent to participate

This study was conducted in compliance with the Ethical Guidelines for Medical and Biological Research Involving Human Subjects and was approved by the Institutional Ethical Committee of Kazusa DNA Research Institute (Approval No. 2020-06).

### Plasma and serum preparation

Blood was collected into EDTA-2K tubes (Venoject II, cat# VP-DK050K, Terumo) using a butterfly needle (Blood Collection Set + Holder, 21G × 3/4", cat# MN-SVS22BH, Terumo) for plasma preparation. Plasma samples were kept at RT or at 4 °C/on ice for the indicated time points, followed by centrifugation for 10 min at 2,000 x g. Subsequently, samples were divided into single-use aliquots and stored at −80 °C until analysis, with no more than two freeze-thaw cycles.

### Cell culture and protein extraction

HEK293T cells (ECACC, Wiltshire, U.K.) were cultured in a 15-cm dish in Dulbecco’s modified Eagle’s medium (Fujifilm Wako, Osaka, Japan) containing 10% fetal bovine serum (Thermo Fisher Scientific, Waltham, MA, USA) at 37 °C, 5% CO_2_, and 80% confluency. The cells were detached using TrypLE Express (Thermo Fisher Scientific) at 37 °C for 5 min. Cells were collected, washed with PBS, and stored at −80 °C until use.

Proteins in HEK293T cells were extracted using protein extraction buffer (100 mM Tris-HCl, pH 8.0, 20 mM NaCl, and 10% acetonitrile (ACN) containing 4% sodium dodecyl sulfate) via sonication in a Bioruptor II (CosmoBio, Tokyo, Japan) for 15 min. The protein concentration in the extract was determined using a BCA protein assay kit (Thermo Fisher Scientific) per the manufacturer’s instructions and was adjusted to 100 ng/µL using a protein extraction buffer.

### Preparation of plasma

Plasma was treated using the Lycopersicon esculentum lectin (LEL) method as previously described,^14^ with modifications. Initially, 25 µL of streptavidin beads suspension (Cytiva, Marlborough, MA, USA) was added to 600 µL of protein-free blocking buffer (Setsuyaku-Kun Supporter, DRC, Tokyo, Japan) diluted 10-fold with Tris-buffered saline (TBS, 25 mM Tris-HCl pH 7.4, 1.37 mM NaCl, and 2.68 mM KCl). Next, 10 µL of 1 µg/µL LEL (Vector Laboratories, Burlingame, CA, USA) was added to the solution. The solution was mixed gently for 30 min and the beads were washed once with 1.2 mL of dilution/wash buffer (TBS with 0.0005% tween20). Next, 25 µL of plasma, diluted in 475 µL of dilution/wash buffer, was added to the beads and mixed for 60 min. Subsequently, the beads were washed twice with 1.2 mL of dilution/wash buffer and mixed in 200 µL of the protein extraction buffer for 15 min. The eluted proteins were then collected. The LEL method was performed automatically using a Maelstrom 9610 instrument (Taiwan Advanced Nanotech, Taoyuan, Taiwan).

The antibody column depletion method used Top14 Abundant Protein Depletion Mini Spin Columns (Thermo Fisher Scientific) per the manufacturer’s instructions (TOP14D method). Plasma samples were pretreated and digested into peptides using the Proteograph workflow (Seer Inc., Redwood City, CA, USA).^15^

### Protein digestion

HEK293T lysate (20 μg) and 200 µL of treated plasma by the LEL method or TOP14D method were subjected to cleaning and digestion with the SP3-LASP method^16^ with minor modifications utilizing the Maelstrom 9610 instrument. The details of SP3-LASP were provided in the Supplemental.

### Dissolution of commercial human cell digests

HeLa cell tryptic digest (Thermo Fisher Scientific; 20 μg) and 100 μg of K562 cell tryptic digest (Promega) were added to 100 μL and 500 μL of 0.02% DMNG containing 0.1% TFA, respectively, and then mixed for 10 min. The concentration of each digest was 25, 50, 100, and 200 ng/μL.

### Preparation of mixed-species samples (human and yeast, *Escherichia coli*) as quantification benchmarks

HeLa cell tryptic digests, yeast digests (Promega, Madison, WI, USA), and *Escherichia coli* digests (Waters, Milford, MA, USA) were dissolved in 0.01% DMNG. HYE mix A contained 65% humans, 15% yeast, and 20% *E. coli*, whereas HYE mix B contained 65% humans, 30% yeast, and 5% *E. coli*. The final mix concentration was 200 µg/µL.

### Nano LC-MS/MS

The tryptic peptides were injected onto a pre-column (PepMap C18, 5 mm × 300 μm × 5 μm, Thermo Fisher Scientific) and separated, at 60 °C, using two analytical columns: 75 μm × 25 cm columns equipped with emitter (Aurora series, CSI, 1.7 µm C18, IonOpticks, Collingwood, VIC, Australia). The separation process was conducted using a NanoElute2 system (Bruker Daltonics, Bremen, Germany) with solvents A (0.1% formic acid in water) and B (0.1% formic acid in acetonitrile). The gradient conditions varied depending on column length and duration.50 min gradient consisted of 0 min 5% B, 2 min 7% B, 42 min 31% B, and 43 min 50% B at a flow rate of 300 nL/min and 43.5 min 70% B and 50 min 70% B at a flow rate of 350 nL/min. 100 min gradient consisted of 0 min 5% B, 92 min 31% B and 93 min 50% B at a flow rate of 300 nL/min and 93.5 min 70% B and 100 min 70% B at a flow rate of 350 nL/min. The peptides eluted from the column were analyzed on a TimsTOF HT (Bruker Daltonics) via Captive Spray II (Bruker Daltonics) and measured in diaPASEF mode using timsControl 5.0. The source parameters were as follows: capillary voltage, 1,600 V; dry gas, 3.0 L/min; and dry temperature, 180 °C. The MS1 and MS2 spectra were collected in the *m/z* range of 100–1,700. The collision energy was set by linear interpolation between 59 eV at an inverse reduced mobility (1/K_0_) of 1.60 versus/cm2 and 20 eV at 0.6 versus/cm2. To calibrate ion mobility dimensions, two ions from the Agilent ESI-Low Tuning Mix were selected (*m/z* [Th], 1/K_0_ [Th]: 622.0289, 0.9848; 922.0097, 1.1895).

For the investigation of the 1/K_0_ range, the Default-diaPASEF method was based on Bruker’s application method for long gradients, with a 1/K_0_ range of 0.6–1.6 and an *m/z* range of 400– 1,200. Using this Default-diaPASEF method as a reference, a new diaPASEF method with a 1/K_0_ range of 0.7–1.3 was designed, referred to as the 0.7–1.3-diaPASEF method (Fig. S1A). Both the Default-diaPASEF and 0.7–1.3-diaPASEF methods were configured with a ramp time of 100 ms, an isolation window width of 26 Th, and a 1 Th overlap for the isolation window. To evaluate the polygon size, the 0.7–1.3-diaPASEF method (ramp time: 100 ms, isolation window: 26 Th, 1 Th overlap) was used as a reference. Based on the 0.7–1.3 diaPASEF method, diaPASEF methods with split isolation windows were generated, specifically in 2-split and 3-split configurations (Fig. S1B). Three additional diaPASEF scans, Narrow1-diaPASEF, Narrow2-diaPASEF, and Narrow3-diaPASEF, were designed by gradually narrowing the polygon size and focusing on regions with high precursor density (Fig. S2A). For variation in ramp time, three diaPASEF methods were created based on Narrow2-diaPASEF, with ramp times set to 100, 150 and 200 ms, respectively (Fig. S2B). To assess the isolation window width, the Narrow2-diaPASEF method (ramp time: 150 ms, 1 Th overlap) was employed and isolation window widths of 16, 26, 36, and 46 Th were tested (Fig. S2C). Based on the evaluation results, the optimal parameters were determined as follows: a Narrow2 polygon size, ramp time of 150 ms, isolation window width of 26 Th, and 1 Th overlap. The optimized diaPASEF method was designated the Thin-diaPASEF method. For comparison, the Thin-diaPASEF method was evaluated along with the py-diaAID PASEF, Slice-PASEF-1F, Slice-PASEF-4F, and Synchro-PASEF methods. The details of each method are provided in the Supplemental Information.

## Data analysis

The DIA-MS files were searched against an *in silico* human spectral library using DIA-NN (version 1.9.2, https://github.com/vdemichev/DiaNN).^17^ First, a spectral library was generated from the UniProt human protein sequence database (downloaded 2024, March, 20,575entry, UP0000056), yeast protein sequence database (downloaded 2024, March, 6,060 entry, UP000002311), *E. coli* protein sequence database (downloaded 2024, March, 4,403 entry, UP000000625), and *C. elegans* protein sequence database (downloaded 2024, July, 19,832 entry, UP000001940) using DIA-NN. To optimize the diaPASEF parameters, the human protein database and spectral library were utilized. To evaluate quantitative accuracy, a protein database and a spectral library comprising three genera, human, yeast, and *E. coli*, were employed.

Additionally, entrapment false discovery rate (FDR) estimation was performed using the Human and *C. elegans* protein databases, along with the corresponding spectral library. The parameters for generating the spectral library were as follows: digestion enzyme, trypsin; missed cleavage, 1; peptide length range, 7–45; precursor charge range, 2–4; precursor *m/z* range, 400–1,200; and fragment ion *m/z* range, 200–1,800. “FASTA digest for library-free search/library generation;” “deep learning-based spectra, RTs, and IMs prediction;” “n-term M excision;” and “C carbamidomethylation” were enabled. The DIA-NN (version 1.9.2) search parameters were as follows: mass accuracy, 15 ppm; MS1 accuracy, 15 ppm; protein inference and gene expression. The MBR was turned off. The protein identification threshold was set at 1% or less for both precursor and protein FDRs.

A protein containing this unique peptide was selected for further analysis. For quantification analysis using mixed species (human, yeast, and *E. coli*), proteins were selected from valid values detected in at least 70% of the samples within at least one experimental group. The coefficient values were calculated using Perseus v1.6.15.0 (https://maxquant.net/perseus/).^18^

For the entrapment method, LC-MS data were analyzed using DIA-NN (version 1.9.2), Spectronaut v19,^19^ and PaSESR 2023b, a mixed uniport human and *C. elegans* protein database.

For DIA-NN, an in silico spectral library was initially generated from human and *C. elegans* mixed protein databases, and analysis was performed using the same parameters as those employed to optimize diaPASEF parameters. Spectronaut was used in directDIA+ mode. The analysis parameters were digestion enzyme, trypsin; missed cleavage, 2; peptide length range, 7-52; precursor charge range, 2-4; modifications, "Carbamidomethylation (C),” "Acetyl (Protein N-term)" and "Oxidation (M)"; and MS1 and MS2 tolerance, set to 15 ppm. For PaSER 2023b, an in silico spectral library was generated using DIA-NN (version 1.8.1) and analyzed. The protein identification threshold was set to 1% or less. As protein inference does not work well in PaSER 2023b, proteins with at least one unique peptide, besides the FDR filter, were selected. To illustrate the overlap of the identified proteins among the different pretreatment methods used for plasma proteome analysis, an UpSet plot was generated using Python.

## Result and discussion

### Optimization of the diaPASEF method for deep proteomics

To optimize the diaPASEF method, several parameters were systematically evaluated to enhance its performance. The key parameters included the collisional cross-section (CCS) dimensions, isolation window division, origin region, ramp time, and isolation window width. Each parameter was assessed using HEK293 digestion with five iterations per condition to ensure reproducibility.

Initially, the CCS dimensions were examined. In Bruker’s application method for long gradients, the default diaPASEF has a 1/K_0_ range of 0.6–1.6. In contrast, the *m/z* range generally analyzed by DIA is around 400–1200 *m/z*, and a 1/K_0_ range of 0.7–1.3 is sufficient. Because ions accumulate in the TIMS device and are subsequently separated in the CCS, limiting the 1/K_0_ range is expected to increase the enrichment in the CCS, which in turn is expected to enhance the intensity and separation of the precursor. Therefore, we analyzed the diaPASEF method using the Default-diaPASEF with a 1/K_0_ range of 0.6–1.6 and compared it with the diaPASEF method having a 1/K_0_ range of 0.7–1.3 (0.7–1.3-diaPASEF). The 0.7–1.3-diaPASEF method also increased protein and precursor identities. Additionally, the number of proteins with a coefficient of variation (CV) less than 10% increased by approximately 1.15-fold using the 0.7–1.3-diaPASEF method, and the number of precursors increased by approximately 2-fold. This increase was attributed to the limited 1/K_0_ range, which enhanced precursor enrichment efficiency. As a result, the precursor intensity increased, improving protein and peptide identity and enabling more stable detection.^20^ In practice, the precursor intensity was approximately 1.5 times higher in the 0.7–1.3-diaPASEF method than that in the Default-diaPASEF method (Fig. S3). Therefore, for further studies, we optimized the conditions using the 0.7–1.3-diaPASEF method (Default2-diaPASEF).

Next, we investigated the potential benefits of splitting the isolation window of diaPASEFs in the 1/K_0_ direction to reduce the number of precursors within a single isolation window. Based on the Default2-diaPASEF method, we created diaPASEF methods in the TimsControl software by splitting the isolation window in the 1/K_0_ direction into two splits (2-split diaPASEF) or three splits (3-split diaPASEF). The cycle times for Default2-diaPASEF and 2-split diaPASEF were 1.91 s, while the cycle time for 3-split diaPASEF was 2.44 s. As the number of splits increased, the number of identified proteins and precursors decreased. The Default2-diaPASEF method detected the highest number of proteins and precursors. Additionally, Default2-diaPASEF identified most proteins and precursors with a CV of less than 10%. However, increasing the number of splits did not provide any benefits; therefore, we continued the optimization using the Default2-diaPASEF method.

Except for some modified peptides, tryptic peptides were concentrated in a specific distribution, allowing the identification of many peptides from this region. Therefore, we aimed to align the polygon region with the concentrated region of tryptic peptides and narrow it to divide the multiple isolation windows in the 1/K_0_ direction. This approach aims to obtain more peptides per PASEF, increase the number of isolation windows, and shorten the cycle time. We created and validated Default2-diaPASEF and three narrow-polygon diaPASEF methods (Fig. S2A). The cycle times for Default2-diaPASEF, Narrow1, Narrow2, and Narrow3 were 1.91 s, 1.56 s, 1.06 s, and 0.74 s, respectively. Narrow2 identified the highest number of proteins and precursors. Interestingly, although Default2-diaPASEF encompassed the widest polygon region, the number of precursors remained almost unchanged until Narrow2. However, Narrow3, which had the narrowest polygon region and shortest cycle time, was too narrow to effectively capture the trace precursors. Narrow2 was selected for further validation because the proportions of proteins and precursors with CV values less than 10% and 20%, respectively, were comparable to those observed in default2-diaPASEF.

Optimization of the ramp time against Narrow1, a parameter directly influencing the peak signal enhancement and cycle time, was conducted. The diaPASEF was conducted with a 100% duty cycle, and three diaPASEF methods were analyzed, featuring Ramp times of 100 ms, 150 ms, and 200 ms, which corresponded to cycle times of 1.06 s, 1.56 s, and 2.06 s, respectively. The diaPASEF method with a ramp time of 150 ms yielded the highest number of identified proteins.

Although the 200 ms Ramp time resulted in a decrease in precursor identification, the number of protein identifications remained comparable to that observed with the 150 ms Ramp time. This may be attributed to precursor saturation within the clusters, whereas an extended accumulation time of 200 ms allowed for the detection of trace proteins. Furthermore, the ratio of the CV values for the proteins and precursors remained consistent across the ramp times. A ramp time of 150 ms was determined to be optimal for maximizing protein and precursor identification.

Finally, the isolation window width was examined. As the width of the isolation window narrowed, the complexity of the MS2 peaks reduced, and peptides were easier to identify; however, the cycle time increased. The Narrow2-diaPASEF method, characterized by a ramp time of 150 ms, was employed with isolation window widths of 15, 25, 35, and 45 Th. These settings corresponded to cycle times of 2.50 s, 1.56 s, 1.25 s, and 1.00 s, respectively. The number of proteins identified increased as the number of proteins narrowed from 45 Th to 25 Th settings, but was slightly less in the 15 Th setting than in the 25 Th setting. This indicates that sufficient relaxation of the MS2 peak was achieved with the 25 Th setting. In addition, the CV for the precursors in the 15 Th settings was approximately 60%, which was approximately 10% lower than those observed for the other isolation window widths. We speculated that the decrease in reproducibility with this 15 Th setting was due to a decrease in the number of data points owing to the extended cycle time. Based on these findings, the 25 Th isolation window width was determined as optimal.

Recent developments in DIA-based proteomics have introduced narrow isolation window widths, such as the 2 Th window used in Orbitrap Astral MS^1^ and the 4 Th window employed in Exploris 480.^4^ These narrower isolation windows allow for the spectral separation of multiple precursors, enhancing specificity and resolution. In contrast, the Thin-diaPASEF method utilizes an isolation window width of 25 Th, which is 12.5 times wider than the 2 Th isolation window of the Astral. However, with the TimsTOF platform, the incorporation of ion mobility separation within the TIMS device enables precursor differentiation not only by *m/z* but also by CCS, effectively increasing the analytical peak capacity 10-fold compared to conventional MS-only separations.^21^ Furthermore, the reduction in spectral complexity achieved through ion mobility separation has been reported to correspond to a fourfold reduction in the effective separation window width.^22^ Despite the relatively wide isolation window of 25 Th used in Thin-diaPASEF, the effective separation corresponds to an approximate isolation window width of 6.5 Th, which ensures sufficient spectral resolution and detection power.

The diaPASEF method with a Narrow1 polygon region and a ramp time of 150 ms, optimized thus far, is named Thin-diaPASEF. Currently, py-diAID PASEF and slice-PASEF are available across various diaPASEF method models, whereas Synchro-PASEF is compatible with the TimsTOF HT, TimsTOF Ultra, and newer models. The Thin-diaPASEF method can be easily implemented using a conventional control software, such as TimsControl, making it compatible with all TimsTOF series instruments. To evaluate its performance, we compared Thin-diaPASEF with other diaPASEF methods, including py-diAID PASEF, Slice-PASEF (Slice-PASEF-1F and Slice-PASEF-4F), and synchro-PASEF. For this comparison, 100 ng of the HKE293 cell digest was analyzed in five replicates, using a 50-min gradient for each method (Fig. 2A). Thin-diaPASEF, py-diAID PASEF, Synchro-PASEF, Slice-PASEF-4F, and Slice-PASEF-1F had the highest number of proteins, in descending order, with average identities of 8,734, 8,636, 8,225, 7,958, and 7,748, respectively. For precursor identification, py-diAID PASEF, Thin-diaPASEF, Synchro-PASEF, Slice-PASEF 4F, and Slice-PASEF 1F averaged 136,212, 126,918, 94,156, 88,258, and 82,638, respectively. Although we expected that the number of precursors in Thin-diaPASEF would be smaller than that of the others because of the narrower polygon region, the number of precursors was the second largest. This indicated that smaller precursor peaks were detected and identified in the selected narrow polygon region. The narrow polygonal area of the thin diaPASEF shortened the cycle time, thereby extending the ramp time. We believe that this prolongation allows the ions to be concentrated and smaller precursor peaks to be detected and identified. By identifying this small precursor, Thin-diaPASEFh was found to have the highest number of proteins, surpassing py-diAID PASEF, which had the highest number of precursors. Furthermore, 95% of the number of proteins identified in Thin-diaPASEF had a CV value of 20% or less, confirming its high reproducibility. These results indicate that Thin-diaPASEF provides higher proteome coverage and robust analysis than the other methods. However, the best reproducibility was achieved with Slice-PASEF-1F at both the protein and precursor levels. Cycle times for each method were 1.5 s for Thin-diaPASEF, 2.2 s for py-diAID PASEF, 0.11 s for Slice-PASEF 1F, 0.64 s for Slice-PASEF-4F, and 0.53 s for Synchro-PASEF, Slice-PASEF-1F was by far the shortest. This shorter cycle time allowed Slice-PASEF-1F to acquire more data points and achieve higher analytical reproducibility. Slice-PASEF-1F is an attractive method for studies requiring high reproducibility, although the number of proteins that can be identified is low. Thus, other methods may have their own suitable uses, and we do not believe that Thin-diaPASEF is the best method for everything; however, we believe that it has an advantage in deep proteome analysis.

**Figure 1.**
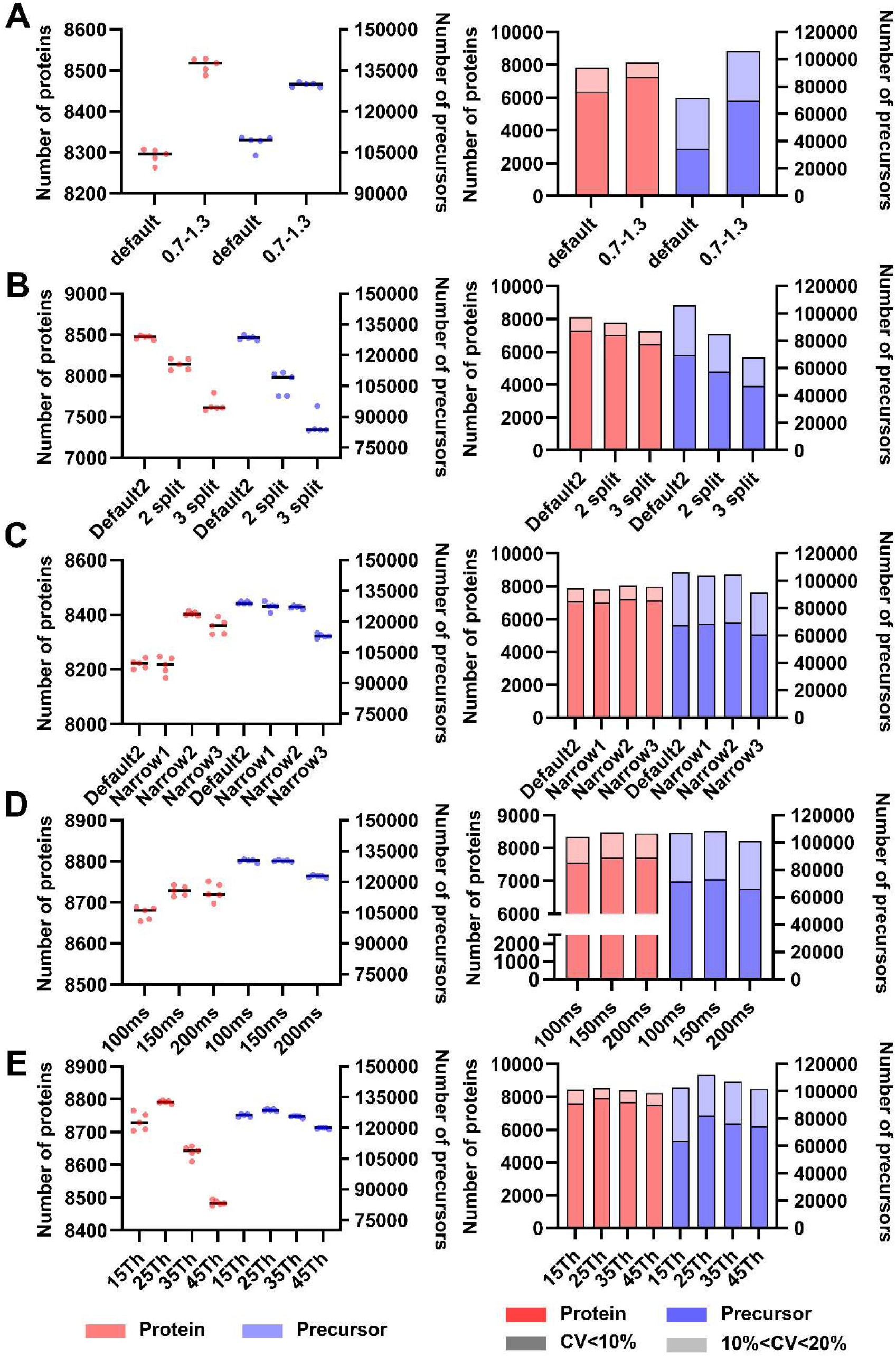
Optimization of the diaPASEF Parameters. Each parameter was evaluated using five replicate measurements of 100 ng digested HEK293 cells. Protein identification consistency and CV values were calculated. A) Impact of narrowing the A 1/K₀ range. A diaPASEF method with a 1/K₀ range of 0.7–1.3 was established. B) Effect of split isolation window. The diaPASEF method with a 1/K₀ range of 0.7–1.3 was named default2-diaPASEF. The isolation window in the Default2-diaPASEF method was further divided into two or three segments for comparison. C) Optimization of polygon size. Starting with the default diaPASEF method, the scan region was progressively narrowed to generate the three diaPASEF methods. D) Effect of ramp time. The ramp time was adjusted to 100, 150, and 200 ms based on the optimized Narrow2 polygon region from (C). E) Optimization of isolation window width. Using the optimized Narrow2 polygon region from (C) and a ramp time of 150 ms from (D), the effects of isolation window widths of 15, 25, 35, and 45 Th were assessed.

**Fig. 2.**
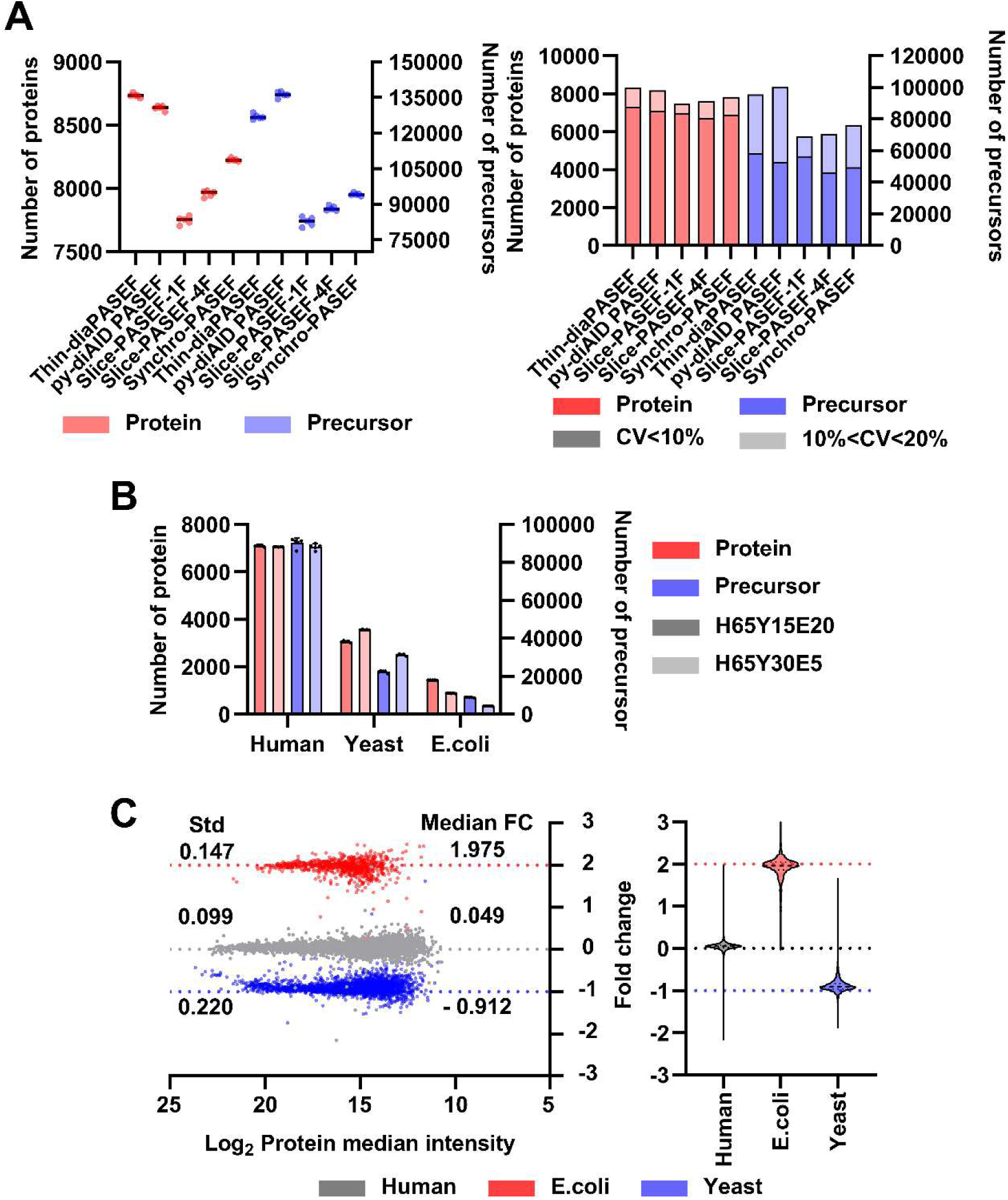
Performance evaluation of Thin-diaPASEF. A) Comparison of DiaPASEF methods. Based on the optimization in Fig. 1, the CCS range was set to 0.7–1.3, the polygon size to Narrow2, the ramp time to 150 ms, and the isolation window width to 25 Th. This optimized diaPASEF method was designated thin-diaPASEF. Five different methods— Thin-diaPASEF, py-diaAID PASEF, slice-PASEF 1F, slice-PASEF 4F, and synchro-PASEF—were evaluated using 100 ng of digested HEK293 cells, and their performances were compared based on the number of identified proteins and CV values for reproducibility. B) Protein identification in mixed-species samples. Two different ratios of human, yeast, and *E. coli* proteins (H65Y15E20: Human 65%, Yeast 15%, *E. coli* 20%; H65Y30E5: Human 65%, Yeast 30%, and *E. coli* 5%) were prepared. Each sample (200 ng of digested peptides) was analyzed in five replicates using Thin-diaPASEF. C) Quantification accuracy of the thin diaPASEFs For quantitative evaluation, proteins quantified in at least 70% of the replicates within each group were selected, and their relative abundance ratios were calculated. The expected ratios for each species were 1, 0.5, and 2 for human, yeast, and *E. coli*, respectively.

### Evaluation of quantitative accuracy using Thin-diaPASEF

To evaluate the quantitative accuracy of each analytical method, we analyzed tryptic digests of human, yeast, and *E. coli* proteins in two distinct mixtures (H65Y15E20:65% human, 15% yeast, and 20% E. coli; H65Y30E5:65% human, 30% yeast, and 5% *E. coli*),^23^ and each mixture was analyzed in five replicates. Protein identifications for human, yeast, and *E. coli* averaged 7,102, 3,059, and 1,437 proteins, respectively, for H65Y15E20, and 7,049, 3,554, and 878 proteins, respectively, for H65Y30E5. Precursor identification averaged 90,206, 22,391, and 8,819 for H65Y15E20, and 88,210, 31,232, and 4,263 for H65Y30E5 (Fig. 2B). Proteins with quantitative values detected in all four replicates for each condition were used for the quantitative analysis. The numbers of Human, Yeast, and *E. coli* proteins used for comparison were 6,812, 2,911, and 812, respectively. The ratios for each organism closely matched the expected ratios (Fig. 2C), and the standard deviations of these ratios were within 0.3, suggesting that quantitative analysis was performed with high accuracy and reproducibility.

### Toward deeper proteome analysis using Thin-diaPASEF

We doubled the gradient time to 100 min for higher-depth analysis using Thin-diaPASEF (Fig. 3A). Similar to the 50 min gradient, analysis of 100 ng of HEK digests in a 100 min gradient did not show a large increase in protein. However, a marked increase was observed starting at 200 ng, with an average of 9,236 proteins identified, and a maximum at 500 ng, with an average of 9,453 proteins identified (Fig. 3A).

**Fig. 3.**
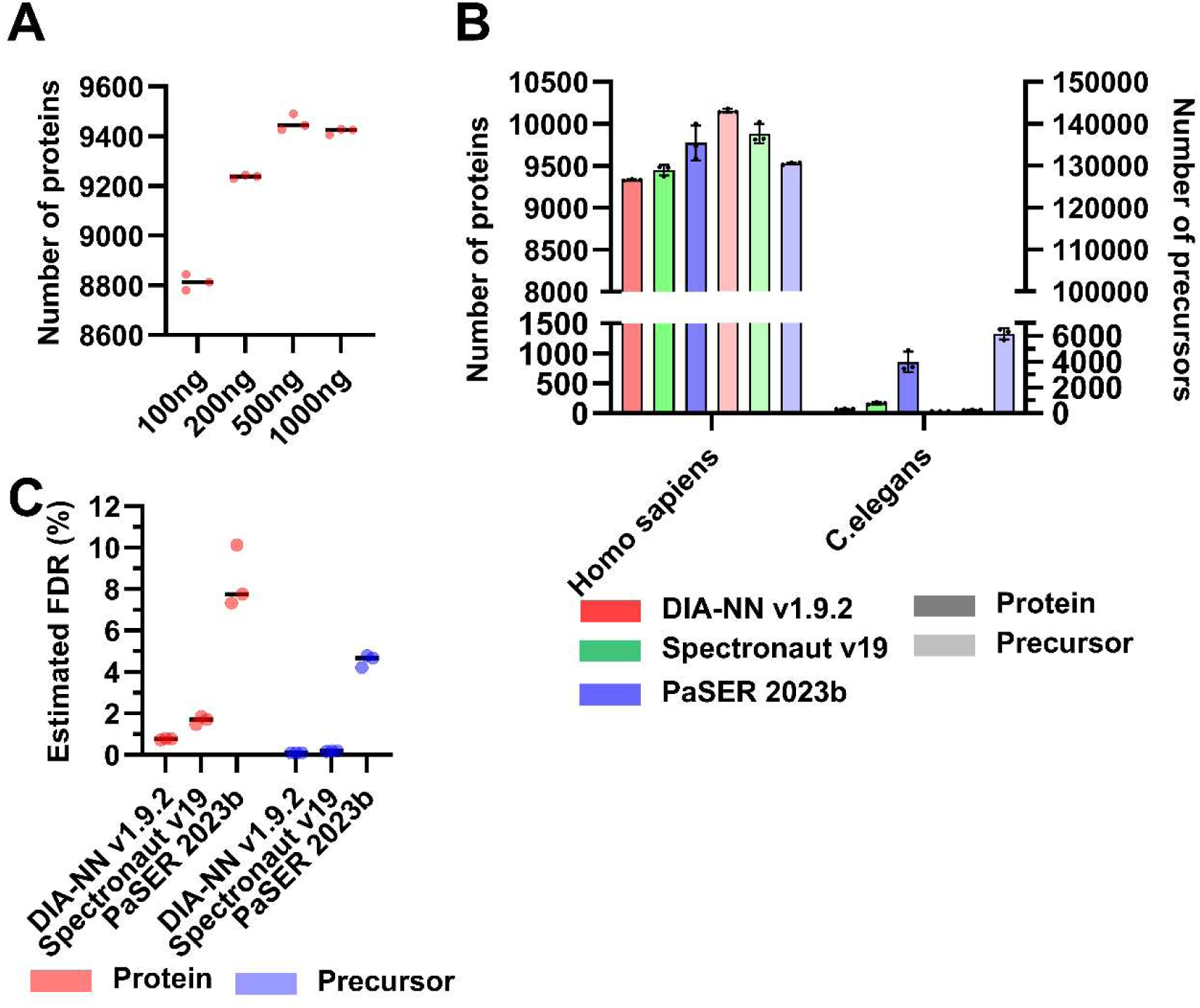
Estimation of false discovery rate (FDR) in maximizing protein identifications. A) Evaluation of peptide loading amounts using a 100 min gradient. To achieve deep proteomics using Thin-diaPASEF, peptide loadings of 100, 200, 500, and 1000 ng were analyzed. B) Identification results for humans and *C. elegans*. The number of proteins and precursors identified in humans and *C. elegans* using DIA-NN, Spectronaut, and PaSER. C) FDR estimation using software tools. Data from the 500 ng peptide load were analyzed using Human and *C. elegans* databases with DIA-NN, Spectronaut, and PaSER to estimate the FDR.

The protein identification analysis software performance is important for in-depth proteomic analysis. In recent years, DIA has enabled the detection and analysis of numerous proteins. However, this advancement also underscores the importance of ensuring the reliability of theFDRs.^24^ To assess the impact of software choice on the reliability of protein identification and quantification in DIA, we compared the performance of PaSER 2023b with TimsDIA-NN, Spectronaut v19,^19^ and DIA-NN v1.9.2^17^ for controlling FDRs. The FDR was estimated using the entrapment method,^25^ which involves adding a non-target species database to the search space to calculate FDRs quasi-statically. Each software tool analyzed the data independently using this approach for each diaPASEF method and cell line using a protein database combined with human and *C. elegans* proteins. Protein inference was ineffective in PaSER 2023b; therefore, proteins quantified using at least one unique peptide were selected for further analysis. Human protein identification increased in the following order: DIA-NN > Spectronaut > PaSER. In the PaSER, some measurements exceeded 10,000 proteins, with an average of approximately 9,750 identified proteins (Fig. 3B). However, PaSER also identified a significant number of C. elegans proteins, raising concerns regarding their specificity. FDR estimates were calculated for these proteins (Fig. 3C). DIA-NN achieved a protein FDR of approximately 0.76%, whereas Spectronaut showed an FDR of approximately 1.64%. In contrast, PaSER exhibited significantly higher protein FDRs, (8.41%, suggesting a lower level of reliability in its identification (Fig. 3B). Only DIA-NN successfully maintained an FDR of < 1%. Based on these results, DIA-NN v1.9.2 was determined to be the most reliable and optimal choice for protein identification. All MS data examined for the methods were analyzed with DIA-NN v1.9.2, and our best protein identification results, with an average of 9,453 proteins, also confirmed that less than 1% of protein FDR controls were valid.

This is the first report of over 9,000 proteins from a single cell type detected in a single-shot analysis in a timsTOF MS series with a correctly controlled protein FDR of less than 1%, proving that the Thin-diaPASEF analysis system is suitable for deep analysis.

### Challenge of plasma proteome analysis

Plasma is a sample with high analytical needs used for biomarker discovery, whereas its wide dynamic range of protein concentration is representative of samples that are difficult to comprehensively analyze using proteome analysis. Recently, however, an amazingly deep plasma proteome analysis was performed in which approximately 5,000 proteins were observed with proper pretreatment and Orbitrap Astral MS.^26,27^ In this study, we attempted to realize a deeper analysis of the plasma proteome by combining an improved version of the LEL method we previously reported, with Thin-diaPASEF measurement using timsTOF HT, without using the expensive Orbitrap Astral MS. In the pretreatment method, Scientific’s top14 abundant protein depletion (TOP14D) method and Seer’s NPA and NPB fractionation were used as controls, in addition to the improved LEL method, followed by enzymatic digestion. Two hundred nanograms of digested samples (200 ng) were analyzed using Thin-diaPASEF with 50-min active gradients, with three replicates for each condition (Fig. 4A). Average of 5,378, 3,465, 2,228, and 1,090 proteins was identified in LEL, NPA, NPB, and TOP14D, respectively. The LEL method detected more proteins than other methods.

**Fig. 4.**
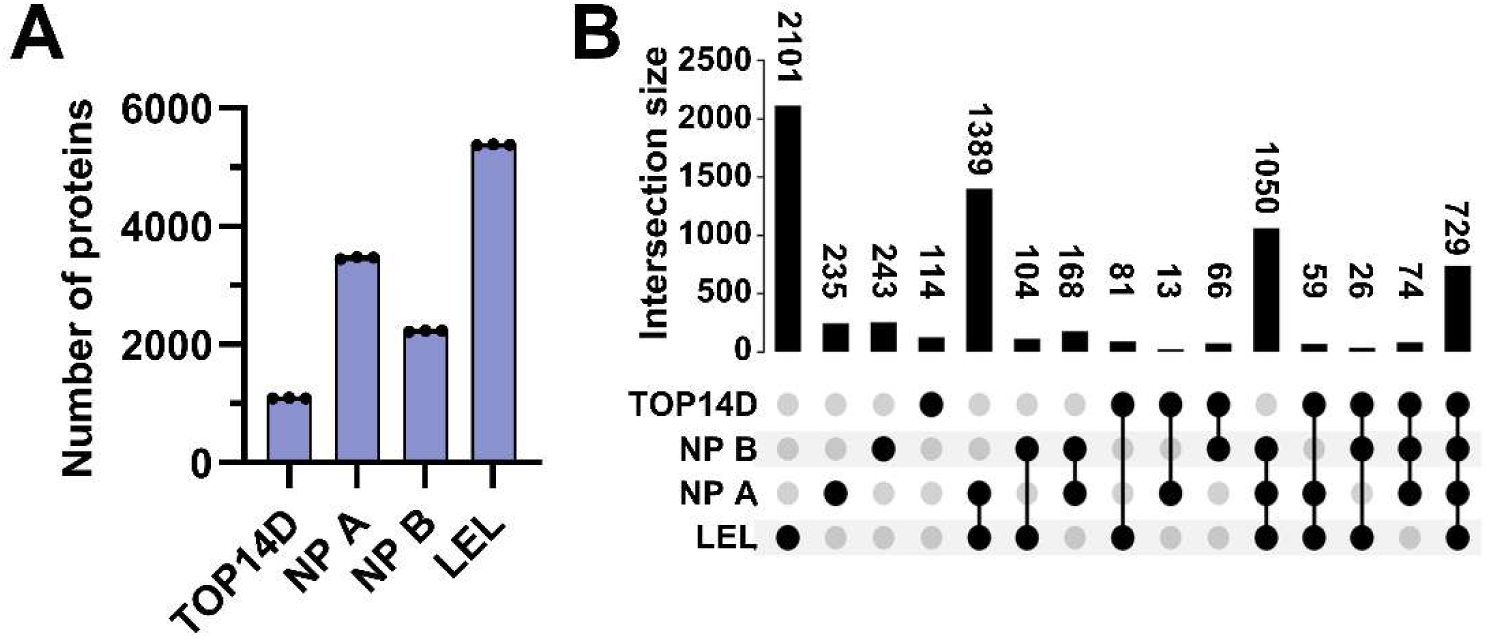
Deep plasma proteome analysis using Thin-diaPASEF. A) Comparison of plasma proteome analysis using the LEL method. Plasma samples were processed using a Top14 removal column (TOP14D method), the LEL method, and Seer’s NPA and NPB nanoparticle-based methods and analyzed using Thin-diaPASEF. Each sample (200 ng) was injected and analyzed in triplicate. B) Overlap of identified proteins across methods. The protein identification results from each preprocessing method were analyzed, and the overlap between them was calculated.

In our previous study, we reported that the combination of *Solanum tuberosum* lectin (STL) and LEL was the best way to enrich low-abundance proteins in serum. However, high-quality biotinylated STL from Vector Laboratories was unavailable for approximately a year (now available). Therefore, in this study, we used LEL, which can enrich trace proteins even when used alone, and optimized the enrichment method. We have previously reported that the STL/LEL method identified approximately 1.5-fold more proteins than the TOP14D method, whereas LEL alone identified more than 4-fold more proteins than those identified in this study. This was mainly due to the optimization of the dilution and wash buffers, with excellent results obtained using LEL alone. In Seer’s NPs technology, which has been the focus of much attention in recent years, NPA has the highest number of proteins identified, with more than 3,000 proteins observed in a single fraction, which is clearly superior to the commonly used TOP14D method. Although the LEL method was superior, the overlap of proteins identified using each method showed that proteins not detected using the LEL method were detected by the other method (Fig. 4B), suggesting that a combination of various methods may be effective for analyzing plasma with higher proteome coverage.

We compared the superior 50-min gradient Thin-diaPASEF combined with the LEL system with previously reported deep plasma proteome analyses. Heil et al. successfully identified 5,163 proteins in samples treated with the Mag-Net method to enrich membrane proteins from plasma using Orbitrap Astral MS with a 60-min gradient.^26^ While this result identified the highest number of proteins reported for plasma proteome analysis using mass spectrometry, to our knowledge, we were able to identify 5,378 proteins in a 50-min gradient using timsTOF HT, which is less expensive than astral. In addition, Beimers et al. measured samples enriched by Seer’s NPA and NPB fractionation methods using Orbitrap Astral MS with a gradient of 30 min each and analyzed the NPA and NPB MS data together to identify approximately 4,600 proteins that have been successfully identified.^27^ In this study, each of the two NPs was measured via LC-MS/MS, so even a 30-min gradient took twice as long, and the identification of 5,378 proteins in a 50-min gradient was again considered an excellent result. This detection level is leading worldwide for deep plasma proteomic analysis, confirming the system’s ability to address comprehensive plasma proteomics. The 50-min gradient was 60 min per measurement, including the overhead time, resulting in an analytical throughput of 24-samples/d. With the ability to observe over 5,000 proteins in the plasma at this throughput, we believe it will be a powerful biomarker discovery system.

## Conclusion

We optimized the diaPASEF method, termed the thin-diaPASEF method, which targets the region with the highest precursor ion concentration in the ion mobility-*m/z* dimension. This approach employs a narrower 1/K_0_ range of 0.7–1.3 than the conventional diaPASEF method’s range of 0.6–1.6. This narrower range enhances the ion mobility separation and effectively concentrates the precursor ions within a specific mobility window. This increased concentration led to a higher precursor intensity, resulting in improved signal-to-noise ratios and enhanced reproducibility across replicate analyses. Furthermore, narrowing the polygon of the diaPASEF window not only enables efficient cycle times but also further enhances precursor intensity and protein identification. Importantly, the Thin-diaPASEF method improves protein identification and achieves high quantitative reproducibility, demonstrating its capability to provide both comprehensive and precise proteomic data. Using a measurement time of 100 min, this optimized method successfully identified approximately 9,400 proteins with a 500 ng sample load. This was the largest number of correctly regulated proteins (FDR <1%) identified from a single cell type in a single-shot analysis of the timsTOF MS series, demonstrating the suitability of our system for deep proteome analysis.

We further applied this optimized Thin-diaPASEF method to plasma proteome analysis. The plasma was processed using an improved LEL method. Remarkably, this approach detected approximately 5,300 proteins in a 50-min measurement (24-samples/d), exceeding the detection capacity and analytical throughput of the best deep plasma proteome system that combines the MagNet method with the expensive Orbitrap Astral MS. Therefore, our system will be highly useful for biomarker discovery.

## Data Availability

Mass spectrometry proteomic data were deposited in the ProteomeXchange Consortium via the jPOST partner repository,^28^ with dataset identifiers PXDxxxxxxx for ProteomeXchange and JPSTxxxxx for jPOST.

## Supporting information

Supplemental infomation

## Acknowledgments

This study was supported in part by JSPS KAKENHI under Grant Numbers 23H02465, 22KK0077, 20K20469, and 21K07877 and by AMED under Grant Number JP22ek0109586. This study was supported by the Kazusa DNA Research Institute. We thank Hiroko Kinoshita, Yukiko Eifuku, Yuka Hayama, Kunihiro Suda, and Chika Takahashi for their help with the reagents, sample preparation, and computational work.

## Author Contributions

YK, RK, and OO conceived and designed the study. MI, DN, and KI prepared the samples. RK conducted LC-MS/MS measurements. RK performed the computational work. YK and RK wrote the manuscript, and MI, DN, KI, and OO edited it. All the authors have read and approved the final version of the manuscript.

## Abbreviations

(ACN): acetonitrile
(DIA): data-independent acquisition
(DMNG): decyl maltose neopentyl glycol
(TIMS): trapped ion mobility spectrometry
(diaPASEF): DIA method using parallel accumulation-serial fragmentation technology
(LC): liquid chromatography
(LEL): *Lycopersicon esculentum* lectin
(STL): *Solanum tuberosum* lectin
(PASEF): parallel accumulation-serial fragmentation
(TFA): trifluoroacetic acid
(TBS): tris buffered saline
(CCS): collisional cross-section
(CV): coefficient of variation

