## Supplemental infomation for "Thin-diaPASEF: diaPASEF for maximizing proteome coverage in single-shot proteomics"

### Supplementary material

Supplementary Method

Supplementary Figure

Supplementary References

### Supplementary Method

Protein digestion using SP3-LASP

Briefly, two types of SeraMag SpeedBead carboxylate-modified magnetic particles (hydrophilic particles, CAT# 45152105050250, and hydrophobic particles, CAT# 65152105050250; Cytiva) were used. These beads were combined in a 1:1 (v/v) ratio, washed twice with distilled water, and reconstituted in distilled water to achieve a concentration of 10 μg solids/μL. Then, 20 μL of the reconstituted beads (SP3-beads) was added to the protein sample, followed by 1-propanol to a final concentration of 75% (v/v), and mixed for 20 min. The beads were collected and washed twice with 80% 1-propanol and once with ethanol. Subsequently, the beads were resuspended in 80 μL of 50 mM Tris-HCl (pH 8.0), 10 mM $CaCl_2$ containing 0.02% lauryl maltose neopentyl glycol, and 2 μL of 500 ng/μL Trypsin Platinum (Promega, Madison, WI, USA). Thereafter, the sample was gently mixed at 37 °C for 14 h to digest the protein. The digested sample was reduced and alkylated with the addition of 8 μL 110 mM tris(2-carboxyethyl)phosphine and 440 mM 2-chloroacetamide at 80 °C for 15 min and then acidified with 16 μL of 5% trifluoroacetic acid (TFA). The sample was desalted using a GL-Tip SDB (GL Sciences, Tokyo, Japan), which was washed with 25 μL of 80% acetonitrile (ACN) in 0.1% TFA, followed by equilibration with 50 μL of 3% ACN in 0.1% TFA. The sample was loaded onto the tip, washed with 80 μL 3% ACN in 0.1% TFA, and eluted with 50 μL of 36% ACN in 0.1% TFA. The eluate was dried using a centrifugal evaporator (miVac Duo Concentrator; Genevac, Ipswich, UK). The dried sample was then redissolved in 0.02% decyl maltose neopentyl glycol (DMNG) containing 0.1% TFA. The peptide concentration in the HEK293T sample was measured using a Pierce Quantitative Fluorescent Peptide Assay kit (Thermo Fisher Scientific, Waltham, MA, USA) per the manufacturer's instructions and adjusted to 100 ng/µL with 0.02% DMNG containing 0.1% TFA.

Comparison of diaPASEF method

The py-diAID PASEF method was generated based on the Thin-diaPASEF method using the Python package for DIA with automated isolation design (py-diAID) software.8 The py-diAID PASEF method was configured according to the parameters reported by Skowronek et.al, with a $1/K_0$ range of 0.7–1.3, ramp time of 100 ms, isolation window width of 25 Th, and other parameters set to default[1]. The Slice-PASEF-1F and Slice-PASEF-4F methods were configured based on the settings reported by Szyrwiel et.al, with a $1/K_0$ range of 0.75–1.2 and a ramp time of 100 ms[2]. The actual polygon regions used in the experiment were downloaded, and their positions were adjusted accordingly. The Synchro-PASEF method was configured per the parameters reported by Skowronek et.al, with a $1/K_0$ range of 0.7–1.3, ramp time of 100 ms, isolation window width of 25 Th, and four synchronized scans[3].

### Supplementary Figure

**A**

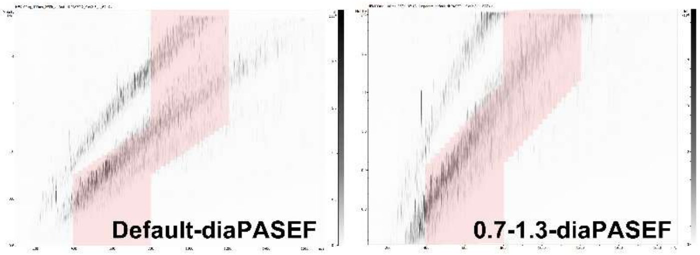


| Method | Default-diaPASEF | 0.7-1.3-diaPASEF |
|---|---|---|
| $1/K_0$ range | 0.6 – 1.6 | 0.7 – 1.3 |
| Ramp time | 100 ms | 100 ms |
| m/z range | *m/z* 400 - 1200 | *m/z* 400 – 1200 |
| Isolation window width | 25 Th | 25 Th |
| Cycle time | 1.91 s | 1.91 s |

**B**

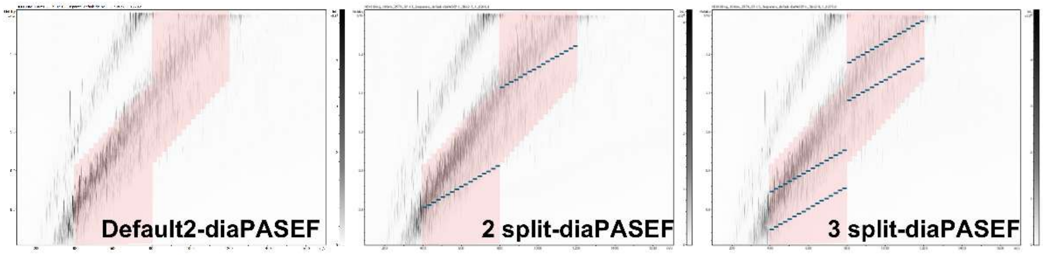


| Method | Default2-diaPASEF | 2 split-diaPASEF | 3 split-diaPASEF |
|---|---|---|---|
| $1/K_0$ range | 0.7 – 1.3 | 0.7 – 1.3 | 0.7 – 1.3 |
| Ramp time | 100 ms | 100 ms | 100 ms |
| m/z range | *m/z* 400 - 1200 | *m/z* 400 – 1200 | *m/z* 400 - 1200 |
| Isolation window width | 25 Th | 25 Th | 25 Th |
| Cycle time | 1.91 s | 1.91 s | 2.44 s |

**Supplementary Fig. S1 Optimized diaPASEF Methods: Narrowed 1/K0 Range and Split Isolation Windows.**

A) Default diaPASEF and narrowed $1/K_0$ range diaPASEF schema. The Default-diaPASEF method embedded in Bruker instruments was based on a $1/K_0$ range of 0.6–1.6. We developed a modified diaPASEF method with a narrowed $1/K_0$ range of 0.7–1.3.

B) Split Isolation Window diaPASEF Schema. Based on the diaPASEF method with a $1/K_0$ range of 0.7–1.3, we created diaPASEF methods with split isolation windows, specifically 2-split and 3-split configurations.

**A**

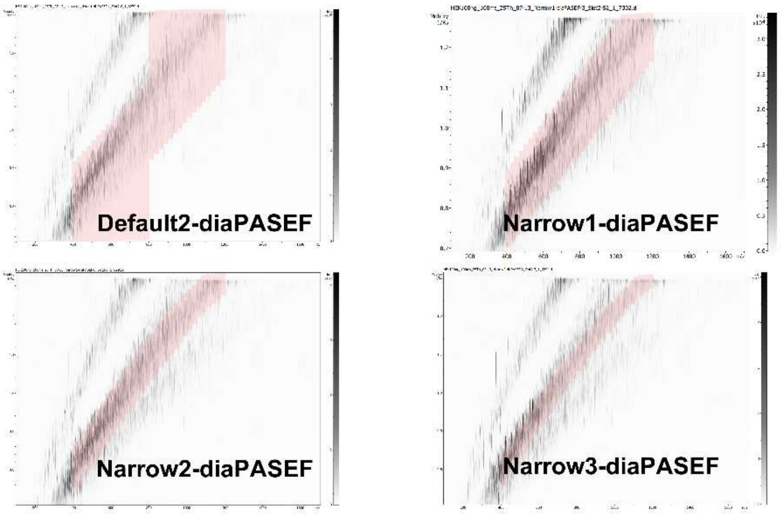


| Method | Default2-diaPASEF | Narrow1-diaPASEF | Narrow2-diaPASEF | Narrow3-diaPASEF |
|---|---|---|---|---|
| $1/K_0$ range | 0.7 – 1.3 | 0.7 – 1.3 | 0.7 – 1.3 | 0.7 – 1.3 |
| Ramp time | 100 ms | 100 ms | 100 ms | 100 ms |
| m/z range | *m/z* 400 - 1200 | *m/z* 400 – 1200 | *m/z* 400 - 1200 | *m/z* 400 - 1200 |
| Isolation window width | 25 Th | 25 Th | 25 Th | 25 Th |
| Cycle time | 1.91 s | 1.06 s | 0.74 s | 0.53 s |

**B**

| Method | 100 ms Narrow1 | 150 ms Narrow1 | 200 ms Narrow1 |
|---|---|---|---|
| $1/K_0$ range | 0.7 – 1.3 | 0.7 – 1.3 | 0.7 – 1.3 |
| Ramp time | 100 ms | 150 ms | 200 ms |
| m/z range | *m/z* 400 - 1200 | *m/z* 400 – 1200 | *m/z* 400 - 1200 |
| Isolation window width | 25 Th | 25 Th | 25 Th |
| Cycle time | 1.06 s | 1.56 s | 2.04 s |

**C**

| Method | 15 Th Narrow1 | 25 Th Narrow1 | 35 Th Narrow1 | 45 Th Narrow1 |
|---|---|---|---|---|
| $1/K_0$ range | 0.7 – 1.3 | 0.7 – 1.3 | 0.7 – 1.3 | 0.7 – 1.3 |
| Ramp time | 150 ms | 150 ms | 150 ms | 150 ms |
| m/z range | *m/z* 400 - 1200 | *m/z* 400 – 1200 | *m/z* 400 - 1200 | *m/z* 400 - 1200 |
| Isolation window width | 15 Th | 25 Th | 35 Th | 45 Th |
| Cycle time | 2.50 s | 1.56 s | 1.25 s | 1.09 s |

**Supplementary Fig. S2 Optimization of diaPASEF Scans: Narrowing Polygon Regions and Varying Ramp Times.**

A) Narrowing Polygon Regions in the diaPASEF scans. Based on the diaPASEF method with a $1/K_0$ range of 0.7–1.3, we focused on regions where precursor ions were concentrated. The polygon regions were progressively narrowed, resulting in the creation of three distinct narrow diaPASEF methods.

B) Variation in Ramp Time for diaPASEF methods. Using the Narrow2-diaPASEF schema as the foundation, we modified the ramp times to 100, 150, and 200 ms.

C) Investigation of the Isolation Window Width. We developed four diaPASEF methods using the Narrow2 schema and a ramp time of 150 ms, varying the isolation window width to 15, 25, 35, and 45 Th.

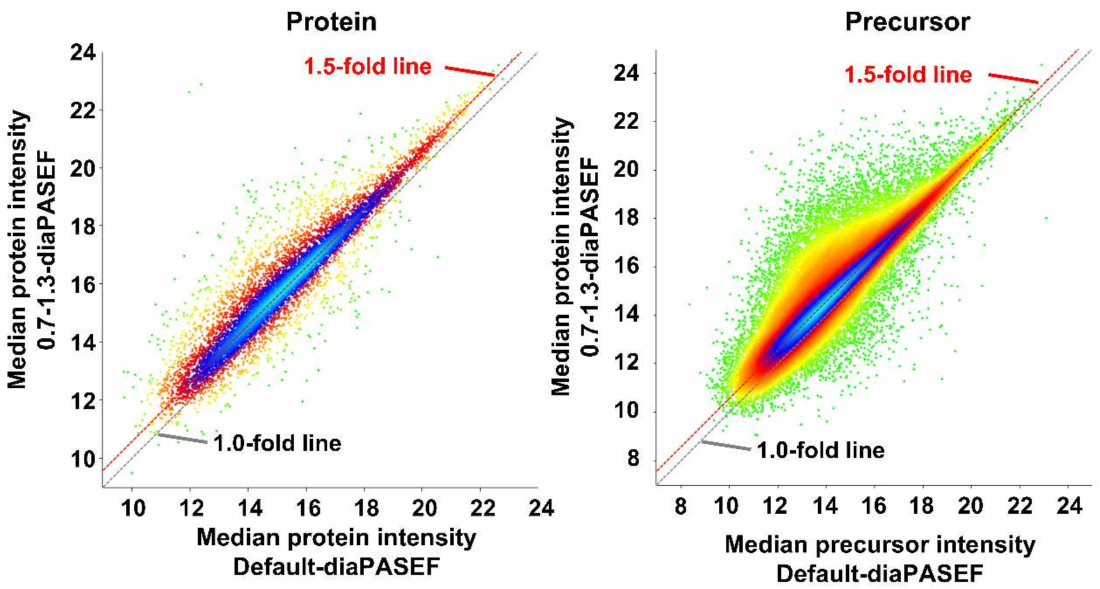


**Supplementary Fig. S3: Intensity Improvement with Narrowed 1/K0 Range**

We calculated and compared the median intensity of quantified proteins and peptides using the Default-diaPASEF
